# Monotreme middle ear is not primitive for Mammalia

**DOI:** 10.1101/2021.05.03.442467

**Authors:** Jin Meng, Fangyuan Mao

## Abstract

The study on evolution of the mammalian middle ear has been fueled by continuous discoveries of Mesozoic fossils in the last two decades. Wang et al.^1^ recently reported a specimen of *Vilevolodon diplomylos* (IMMNH-PV01699)^2^ that adds to the increasing knowledge about the auditory apparatus of ‘haramiyidans’, an extinct Mesozoic group of mammaliaforms. The authors hypothesized that a middle ear with a monotreme-like incus and malleus and incudomallear articulation was primitive for mammals, which challenges the convention that the monotreme middle ear is specialized^3^ or autapomorphic^4^ in mammals. We raise concerns about terminology and identification of the incus presented by Wang et al. and show that their analysis does not support their hypothesis; instead, it supports the one by Mao et al.^5,6^.

---

In their study of “A monotreme-like auditory apparatus in a Middle Jurassic haramiyidan”, Wang et al.^1^ started in promoting a set of terms to replace longstanding ones used for auditory bones with the intention to make them ‘descriptive’. They replaced, for instance, the ‘definitive mammalian middle ear’ (DMME)^7,8^ with ‘detached middle ear’ (DME)^9^. We agree that the DMME is literally imperfect, not because of its descriptive insufficiency but because of its bearing on phylogeny, which has a historical reason. Wang et al. defined the DME as the configuration in which ‘the postdentary bones lack a bony or cartilaginous attachment to the mandible and have an exclusive auditory function’. Compared to the definition for the DMME (“The configuration in which the angular, articular plus prearticular, and quadrate are strictly auditory structures, fully divorced from the feeding apparatus [and renamed tympanic, malleus, and incus].”)^8^, this DME definition is poorly formulated because it does not include the quadrate (incus) and is against the authors’ will to regard the stapes, malleus and incus only as middle ear ossicles. The authors used a mixture of terms including ‘auditory elements’, ‘auditory apparatus’, ‘middle ear’, ‘ossicular chain’, and ‘auditory bones’ in their study, while they disliked ‘auditory bones’ for a good reason. However, using terms such as ‘auditory elements’, as the authors preferred, does not improve the terminology because soft tissues for hearing, such as the stapedius muscle, the tensor tympani muscle, and the tympanic membrane, could be counted in. Moreover, terms such as ‘postdentary attached middle ear’ (PAME) are intrinsically contradictory because the postdentary bones (articular, prearticular, angular, and surangular)^8^ comprise the middle ear, apart from the jaw joint, and are not an attachment tool for the middle ear. ‘Dentary attached’ or ‘postdentary’ middle ear may better describe the configuration in question. However, we caution to keep coining such terms at the minimum while recognizing the history of science and rule of priority. New terms are necessary for novel structures, but it only worsens the already complicated anatomic terminology to replace existing terms with ill-defined new ones.

Wang et al. presented some valuable interpretations on previously known but still poorly understood auditory bones, such as the surangular and ectotympanic, in ‘haramiyidans’. Because these subjects have been extensively treated^2,5,6,10–12^, we focus our discussion on the incus and malleus that Wang et al. provided some new evidence, which in turn led to their conclusion. The authors claimed that in their specimen (IMMNH-PV01699) the ossicular chain is ‘well-preserved and in near-life position’. We note instead that the stapes was never mentioned and that the two sets of ossicular remains were transferred to the left mandible with the incus and malleus from both sides remarkably remaining in articulation. The purported incus and incudomallear articulation in their specimen were described as monotreme-like, but it is notably different from those in the holotype of *V. diplomylos*^2^. The authors also thought that the incus ‘resembles’ and “has a similar outline” to those of the *Jeholbaatar*^13^ and *Arboroharamiya allinhopsoni*^10–12^. However, the ‘incus’ of *Jeholbaatar* has been shown to be part of the malleus by new evidence^6^, as noted by Wang et al. The only known unequivocal incus of euharamiyidans was from *Arboroharamiya,* a sister taxon of *V. diplomylos*, which has been repeatedly described as convex, bulbous, and like that of therians^10–12^. The ‘incus’ in *Qishou* (see ED fig. 6, ref. 1) is a misinterpretation – it is part of the element with debatable identity (Fig. 1k-m). Why is the purported incus in IMMNH-PV01699 so different from its sister-taxon but similar to monotremes? The possibility that it is part of the other element (identified as the malleus), as in *Jeholbaatar* and *Qishou*, cannot be ruled out. If so, it explains why the incus and malleus “were moved to that degree from their position in life and yet remain well preserved”^1^. We suspect whether Wang et al.’s CT data with a voxel size of 32.7μm could secure the identities of the incus and malleus but cannot verify this because the digital data were not yet available.

**Fig. 1.**
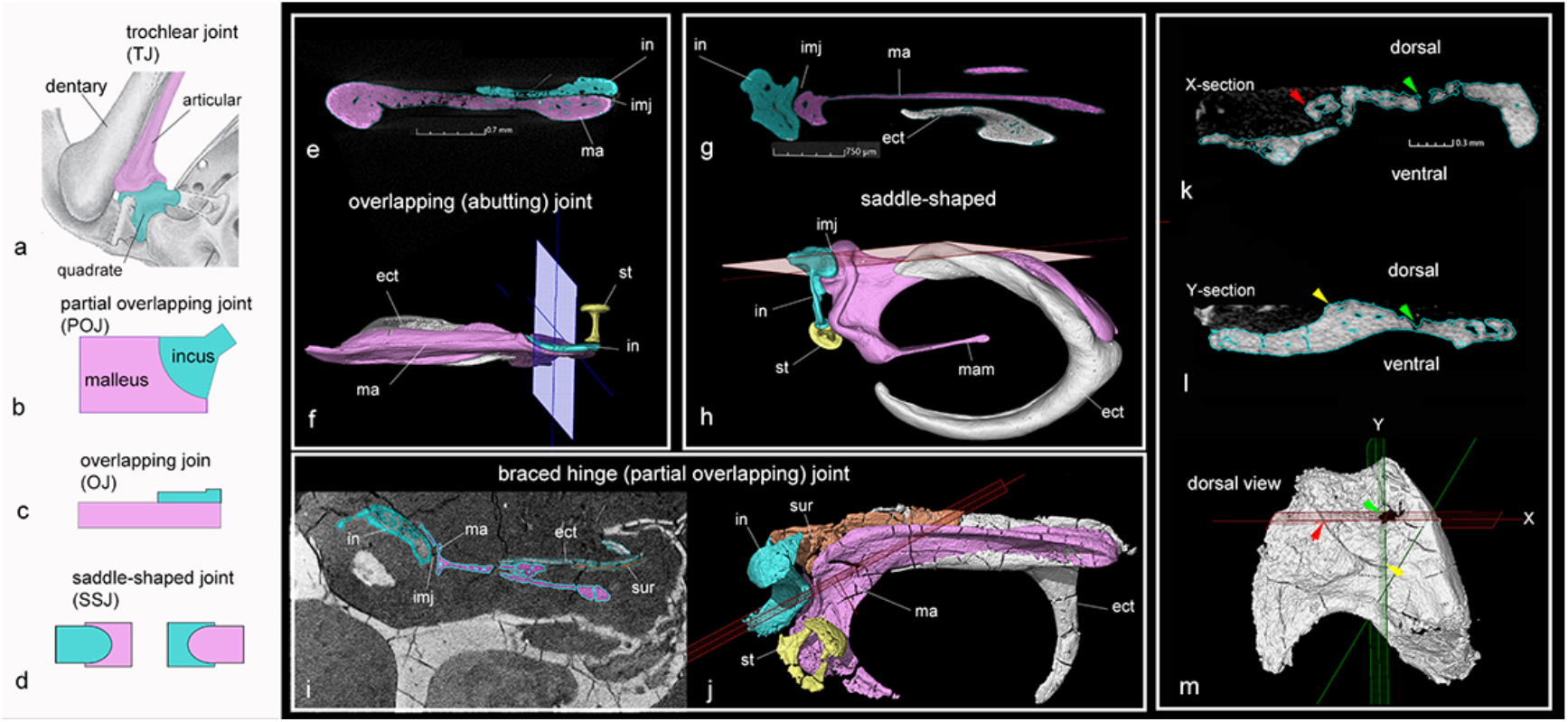
Types of incudomallear articulation. **a**, Mandibular middle ear of *Morganucodon*, showing the trochlear incudomallear joint. **b**, Diagram showing the braced hinge (partial overlapping) joint in cross-sectional view, which is based on **i** and **j**; the incus is largely caudal to the malleus but laterally braced by a bony lip of the malleus^6^. **c**, Diagram showing the overlapping (abutting) joint in cross-sectional view, which is based on **e** and **f**; the incus is dorsal to the malleus. **d**, Diagram showing the saddle-shaped incudomallear joint in lateral (left) and dorsal (right) views, which are based on **g** and **h**, respectively; the incus is caudal to the malleus. **e, f**, A CT-slice (**e**; voxel size of 8.153μm) that runs through the CT rendered ossicles of the monotreme *Tachyglossus* (**f**). **g, h**, A CT-slice (**g**; voxel size of 6.611μm) that runs through the CT rendered ossicles of the marsupial *Didelphis* (**g**); **i, j**, A CT-slice (**i**; voxel size of 7.474μm) that runs through the CT rendered ossicles of the symmetrodontan *Origolestes.* **k-m**, CT-slices (**k** and **l**; voxel size of 10.2μm) that run through the CT rendered element identified as the ectotympanic (**m**) of the euharamiyidan *Qishou*^12,14^; part of which was misinterpreted as the ‘incus’^1^ (colored arrows in **k** and **l** corresponding to the same reference point in **m**). Abbreviations: **ect**, ectotympanic; **imj**, incudomallear joint; **in**, incus; **ma**, malleus; mam, manubrium of the malleus; **st**, stapes. Fig. 1a is based on ref. 8, 1k-m on ref. 12, and others on refs. 5 and 6.

Contrary to the existing hypothesis that the braced hinge joint (=partial overlapping joint, POJ; Fig. 1b, i, j) is potentially primitive for mammals^6^, Wang et al. concluded that optimization of five incudomallear characters in their phylogeny (Fig. 2a) ‘supports the overlapping joint as primitive for Mammalia. The partial overlapping joint is derived from the overlapping joint (and not vice versa) by the caudal shift of the incus with regard to the malleus’. We noted that the five incudomallear characters in *Vilevolodon* and *Qishou* were coded as identical to those of monotremes, but the incus was not preserved in *Qishou*^12,14^(Fig. 1-m) and the so-called malleus is subject to interpretation^1,10,12^. *Sinobaatar* was coded as ‘?’ for the five characters, although its well-preserved malleus and incus^6^ have been illustrated in ref. 1 (fig. 3). The two species of *Arboroharamiya* were coded as having a plate-like incus, which is factually untrue, as mentioned above. Surprisingly, none of the five incudomallear characters showed up as a synapomorphy at any major node (clade) in the consensus tree (see Suppl. Info. ^1^). Nonetheless, the authors managed to optimize them and map the four types of joints (Fig. 1) on the simplified consensus tree as the hypothesis (Fig. 2a).

**Fig. 2.**
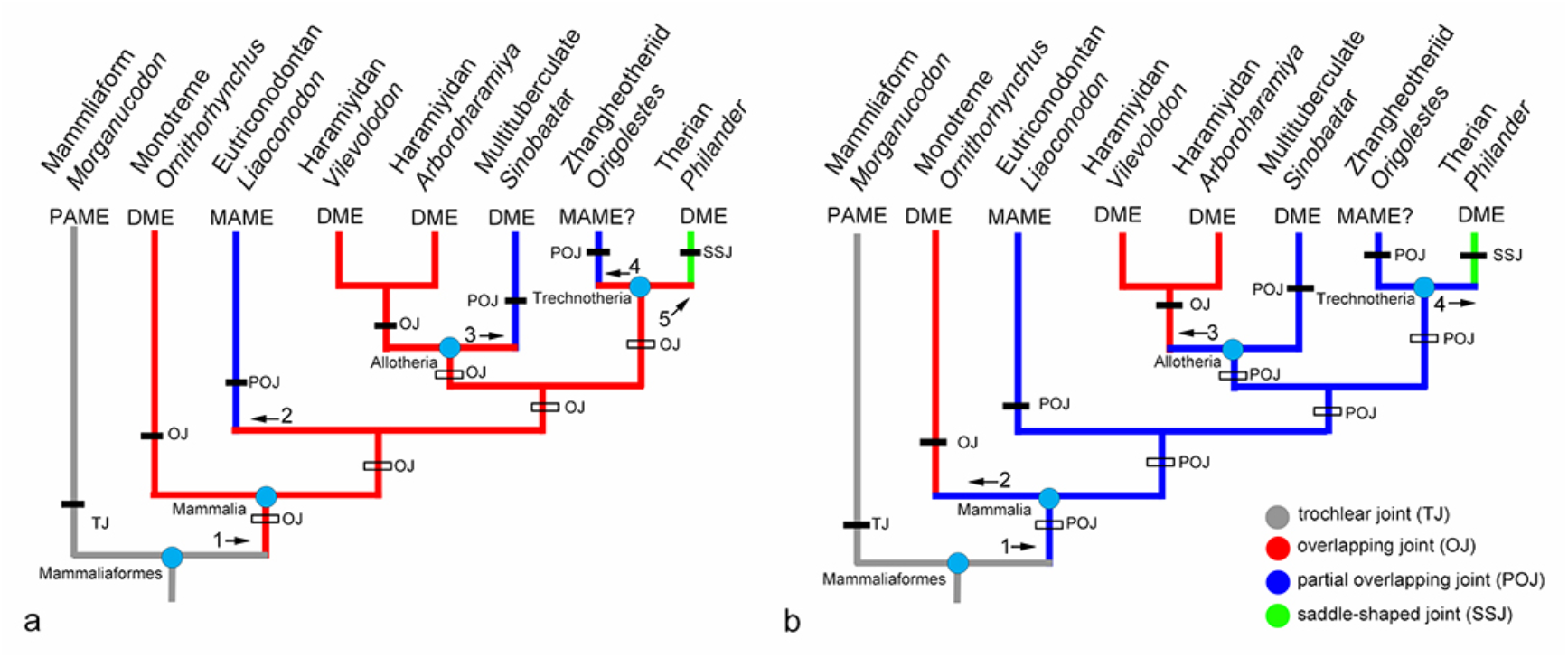
Two hypotheses for evolution of the mammalian middle ear resulted from Wang et al.’s analysis. **a**, Hypothesis preferred by Wang et al.^1^, in which the monotreme-like over-lapping incudomallear joint is primitive for Mammalia. **b**, Alternative hypothesis overlooked by the authors, in which the braced hinge joint (=POJ) is primitive for Mammalia. Taxa, tree topology and optimized character distributions are from the original study (see fig. 3 in ref. 1). We added the empty bars, arrows, and associated labels to visualize the evolutionary changes within the phylogeny. The comparison shows that the overlooked hypothesis b is more parsimonious than a, which supports the existing hypothesis^6^ but rejects the one that monotreme middle ear is primitive for Mammalia.

Under their hypothesis, the first evolutionary step would be from the trochlear joint (TJ) in nonmammalian cynodonts to the OJ of mammals. This step requires several abrupt changes (the transformation through the POJ was deemed impossible by Wang et al.): the incus becoming a flat platelet, complete loss of the synovial joint, and the incudomallear complex transformed to a nearly horizontal position with the incus shifting to the dorsal side of the malleus. As known in developmental studies, the vertical orientation of the ectotympanic in ontogeny was recognized as primitive in mammals^4^ and therians^15^ because the angular bone in nonmammalian cynodonts was vertically positioned. In development of echidna the ectotympanic and malleus perform a ‘flipping’ from their original vertical position to horizontal orientation in adults^16^. The flat incus lying medial to the malleus and a horizontal ectotympanic were considered autapomorphic for monotremes^4^. These studies do not support Wang et al.’s hypothesis. In addition, this step also requires direct change from PAME to DME, another abrupt change supported by no fossil and developmental evidence. Furthermore, within Mammalia, two evolutionary steps from the OJ to POJ took place independently at eutriconodontans and multituberculates and at the node of Trechnotheria, the OJ would give rise either to POJ, which then evolved to the saddle-shaped joint (SSJ) (Fig. 1d, g, h), or to the POJ and SSJ respectively; either evolutionary process involves at least two steps. Thus, a total of at least four evolutionary steps is required within Mammalia.

It appears that Wang et al. have overlooked a better supported result of their optimization and mapping: the POJ is primitive for Mammalia, as we present in Fig. 2b. Under this alternative hypothesis, the evolutionary change from the nonmammalian cynodont OJ to the mammalian POJ would be simple because the incus and malleus retain the trochlear joint, the incus is largely caudal to the malleus, and the auditory bones are positioned nearly vertical; further, they do not need to be fully detached from the dentary. Within Mammalia there are only three evolutionary steps: two independent evolutions of the OJ at monotremes and ‘haramiyidans’, respectively, and one from the POJ to SSJ within Trechnotheria.

Thus, Wang et al. postulated their hypothesis based on the less supported result of their own analysis. Under the rule of parsimony, their hypothesis (Fig. 2a) is falsified because it requires at least five evolutionary steps in the mammalian middle ear evolution. In contrast, their analysis corroborates the alternative (Fig. 2b) that needs only four steps, which supports the existing hypothesis^6^. Wang et al.’s conclusion was likely drawn from a misinterpretation of their result and is therefore misleading. To our knowledge, except for the purported monotreme-like incudomallear joint in IMMNH-PV01699, there is no convincing evidence for such a joint in any non-monotreme mammals and their relatives.

## Author contributions

J.M. and F.M initiated and wrote the paper.

